# Physical exercise increases overall brain oscillatory activity but does not influence inhibitory control in young adults

**DOI:** 10.1101/268995

**Authors:** Luis F. Ciria, Pandelis Perakakis, Antonio Luque-Casado, Daniel Sanabria

**Affiliations:** Mind, Brain, & Behavior Research Center, University of Granada, Spain.; Department of Experimental Psychology, University of Granada, Spain.; Universidad Loyola Andalucía, Departamento de Psicología, Campus de Palmas Altas, 40000, Sevilla, España; Faculty of Physical Activity and Sport Sciences, Pablo de Olavide University, Spain.

**Keywords:** brain rhythms, EEG, information processing, executive control

## Abstract

Extant evidence suggests that acute exercise triggers a tonic power increase in the alpha frequency band at frontal locations, which has been linked to benefits in cognitive function. However, recent literature has questioned such a selective effect on a particular frequency band, indicating a rather overall power increase across the entire frequency spectrum. Moreover, the nature of task-evoked oscillatory brain activity associated to inhibitory control after exercising, and the duration of the exercise effect, are not yet clear. Here, we investigate for the first time steady state oscillatory brain activity during and following an acute bout of aerobic exercise at two different exercise intensities (moderate-to-high and light), by means of a data-driven cluster-based approach to describe the spatio-temporal distribution of exercise-induced effects on brain function without prior assumptions on any frequency range or site of interest. We also assess the transient oscillatory brain activity elicited by stimulus presentation, as well as behavioural performance, in two inhibitory control (flanker) tasks, one performed after a short delay following the physical exercise and another completed after a rest period of 15’ post-exercise to explore the time course of exercise-induced changes on brain function and cognitive performance. The results show that oscillatory brain activity increases during exercise compared to the resting state, and that this increase is higher during the moderate-to-high intensity exercise with respect to the light intensity exercise. In addition, our results show that the global pattern of increased oscillatory brain activity is not specific to any concrete surface localization in slow frequencies, while in faster frequencies this effect is located in parieto-occipital sites. Notably, the exercise-induced increase in oscillatory brain activity disappears immediately after the end of the exercise bout. Neither transient (event-related) oscillatory activity, nor behavioral performance during the flanker tasks following exercise showed significant between-intensity differences. The present findings help elucidate the effect of physical exercise on oscillatory brain activity and challenge previous research suggesting improved inhibitory control following moderate-to-high acute exercise.

## Introduction

Inhibitory control seems to benefit from a previous acute bout of exercising (e.g., cycling or running) at a moderate-to-high intensity (Hillman, Erickson & Kramer, 2008). This effect has been associated with increased neural efficiency as a function of the exercise demands (Erickson, Hillman & Kramer, 2015). However, the neurophysiological pathways by which exercise exerts its beneficial effect on cognition remain unclear. In particular, there is scant evidence on the exact effect of exercise on brain oscillations and their possible mediation in enhancing cognitive performance. To address these issues, here we investigate oscillatory brain activity during an acute 30’ bout of moderate-to-high aerobic exercise, as well as its impact on subsequent inhibitory control.

Previous research on brain dynamics under physical effort has reported a selective tonic power increase in the alpha frequency band in anterior sites (Kubitz & Pothakos, 1997; Boutcher, 1993; Petruzzello et al., 1991), which was linked to the beneficial effects of acute exercise on mood (Lattari et al., 2014; Boutcher, 1993; Petruzzello et al., 1991) and cognitive function (Chang et al., 2015; Dietrich, 2006). Accumulating evidence, however, points to an overall power increase across the entire EEG frequency spectrum (Ciria et al., 2017; Crabbe & Dishman, 2004), which does not seem to be specific to any particular frequency band or brain location. In fact, the potential exercise-induced effect on other EEG frequency bands and scalp localizations remains poorly understood. To our knowledge, no study so far has attempted to adequately address this crucial issue by applying a data-driven cluster-based analysis (bottom-up, without prior assumptions on any frequency range or site of interest).

Regarding exercise-induced changes in event-related brain oscillatory activity, even less is known. To date, the only study investigating this issue (Chang et al., 2015) reported that improved cognitive performance was accompanied by a greater target-evoked decrement of alpha frequency power during a Stroop task. This effect was observed within the first 15 min after the cessation of the exercise (between 50% and 60% of the heart rate reserve) relative to a control (reading) condition in old adults. The authors concluded that acute exercise may provide neural resources for attentional investment and top-down processes to facilitate cognition. These results are in line with previous meta-analyses (Verburgh et al., 2014; Chang et al., 2012; Lambourne & Tomporowski, 2010) pointing to moderate-to-high acute exercise (between the 60% and 80% VO2max) during more that 20 minutes as the key intensity and duration to induce cognitive enhancement (particularly in executive processing) between 10 and 20 minutes after the end of the exercise. However, the study by Chang and collaborators (2015) was restricted to the alpha frequency band at frontal locations. Once again, a stepwise cluster-based analysis will provide novel and complementary information for a deeper understanding of the transient exercise-induced changes in event-related brain oscillatory activity.

The present study was therefore designed to investigate the following open questions: 1) does exercise produce an overall increase of the entire EEG frequency spectrum or is the effect localized at specific frequency bands and electrode sites? 2) does moderate-to-high aerobic exercise exert a positive effect on the behavioral performance at an inhibitory control task delivered after the cessation of the exercise? 3) is this positive effect accompanied by specific transient, event-related modulations of particular brain rhythms? 4) for how long do the exercise-induced cognitive benefits last after the termination of the exercise?

To this aim, we compared the oscillatory brain activity (by means of a stepwise cluster-based analysis) of a set of healthy young adults during two acute bouts of aerobic exercise (cycling) at different intensities (in two separate experimental sessions), corresponding to the 80% and 20% of their ventilatory anaerobic threshold (VAT), during 30 minutes. The 20% condition was included as the light intensity exercise baseline condition (instead of a rest non-exercise condition) to match conditions in terms of movement. Further, to explore the time course of exercise-induced changes on brain function and cognitive performance, we included two inhibitory control (flanker; (Eriksen & Eriksen, 1974) tasks. One was performed within the first 10 to 20 minutes after exercise cessation, where the largest effects of moderate-to-high acute exercise are expected according to previous meta-analytical reviews (Verburgh et al., 2014; Chang et al., 2012; Lambourne & Tomporowski, 2010). The second flanker task was delivered following a 15’ resting period after the first task. We expected a higher power increase across the entire EEG frequency spectrum during moderate-to-high intensity exercise with respect to the light intensity exercise and rest, which would be also accompanied by higher cognitive performance and a distinctive oscillatory brain activity pattern of (task relevant) stimulus processing in the first flanker task. We did not expect significant between-intensity differences in the second flanker task.

## Methods and design

### Participants

We recruited 20 young males (19-32 years old, average age 23.8 years old) from the University of Granada (Spain). All participants met the inclusion criteria of normal or corrected to normal vision, reported no neurological, cardiovascular or musculoskeletal disorders, were taking no medication and reporting less than 3 hours of moderate exercise per week. Participants were required to maintain regular sleep-wake cycle for at least one day before each experimental session and to abstain from stimulating beverages or any intense exercise 24 hours before each session. From the 20 participants, one was excluded from the analyses because he did not attend to the last experimental session and another one because of technical issues. Thus, only data from the remaining 18 participants are reported. All subjects gave written informed consent before the study and received 20 euros for their participation. The protocol was approved in accordance with both the ethical guidelines of the University of Granada and the Declaration of Helsinki of 1964.

### Apparatus and materials

All participants were fitted with a Polar RS800 CX monitor (Polar Electro Öy, Kempele, Finland) to record their heart rate (HR) during the incremental exercise test. We used a ViaSprint 150 P cycle ergometer (Ergoline GmbH, Germany) to induce physical effort and to obtain power values, and a JAEGER Master Screen gas analyser (CareFusion GmbH, Germany) to provide a measure of gas exchange during the effort test. Flanker task stimuli were presented on a 21-inch BENQ screen maintaining a fixed distance of 50 cm between the head of participants and the center of the screen. E-Prime software (Psychology Software Tools, Pittsburgh, PA, USA) was used for stimulus presentation and behavioural data collection.

### VAT determination test

Participants came to the laboratory, one week before the first experimental session to provide the informed consent, complete an anthropometric evaluation (height, weight and body mass index) and to familiarize with the cycle-ergometer and the cognitive task. Subsequently, they performed an incremental cycle-ergometer test to obtain their VAT which was used in the experimental sessions to adjust the exercise intensity individually. The incremental effort test started with a 3 minutes warm-up at 30 Watts (W), with the power output increasing 10 W every minute. Each participant set his preferred cadence (between 60-90 rpm • min-1) during the warm-up period and was asked to maintain this cadence during the entire protocol. The test began at 60 W and was followed by an incremental protocol of 30 W every 3 minutes. Each step of the incremental protocol consisted of 2 minutes of stabilized load and 1 minute of progressive load increase (5 W every 10 seconds). The oxygen uptake (VO_2_ ml • min-1 • kg-1), respiratory exchange ratio (RER; i.e., CO_2_ production • O_2_ consumption-1), relative power output (W • Kg-1) and heart rate (bpm) were continuously recorded throughout the test. VAT is considered to be a sensitive measure for evaluating aerobic fitness and cardiorespiratory endurance performance (Londeree, 1997; Wasserman, 1984). It was defined as the VO_2_ at the time when RER exceeded the cut-off value of 1.0 (Davis et al., 1976; Yeh et al., 1983) and did not drop below that level during the 2 minutes constant load period or during the next load step, never reaching the 1.1 RER (see Luque-Casado et al., 2013; Luque-Casado et al., 2016b, for a similar protocol). The submaximal cardiorespiratory fitness test ended once the VAT was reached.

### Procedure

Participants completed two counterbalanced experimental sessions of approximately 120 min each. Sessions were scheduled on different days allowing a time interval of 48‒72 hours between them to avoid possible fatigue and/or training effects. On each experimental session (see Fig. 1), participants completed a 15’ resting state period sitting in a comfortable chair with closed eyes. Subsequently, they performed 10’ warm-up on a cycle-ergometer at a power load of 20% of their individual VO_2_ VAT, following by 30’ exercise at 80% (moderate-intensity exercise session) or at 20% (light intensity exercise session) of their VO_2_ VAT (see Table 1). Upon completion of the exercise, a 10’ cool down period at 20% VO_2_ VAT of intensity followed. Each participant set his preferred cadence (between 60-90 rpm • min-1) before the warm-up and was asked to maintain this cadence throughout the session in order to match conditions in terms of dual-task demands. Later, participants waited sitting in a comfortable chair until their heart rate returned to within their 130% of heart rate at resting (average waiting time 5’ 44”). The first flanker task was then performed for 6’, followed by a 15’ resting period with closed eyes. Finally, they again completed the 6’ flanker task.

**Figure 1.**
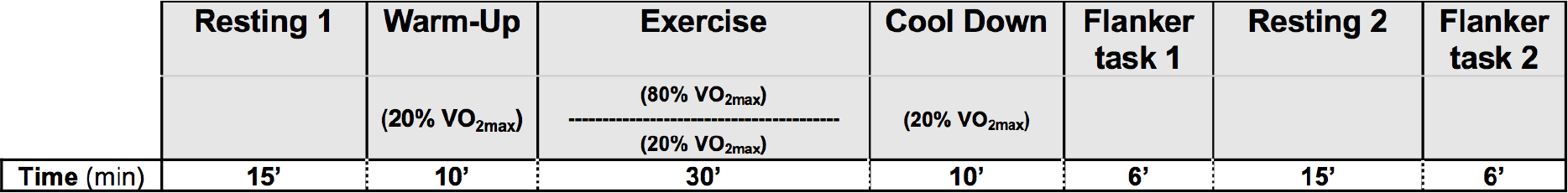
Time course of experimental sessions.

### Flanker task

We used a modified version of the Eriksen flanker task based on that reported in Eriksen and Eriksen (1974). The task consisted of a random presentation of a set arrows flanked by other arrows that faced the same or the opposite direction. In the congruent trials, the central arrow is flanked by arrows in the same direction (e.g., <<<<< or >>>>>), while in the incongruent trials, the central arrow is flanked by arrows in the opposite direction (e.g., <<><< or >><>>). Stimuli were displayed sequentially on the center of the screen on a black background. Each trial started with the presentation of a white fixation cross in a black background with random duration between 1000 and 1500 ms. Then, the stimulus was presented during 150 ms and a variable interstimulus interval (1000‒1500 ms). Participants were instructed to respond by pressing the left tab button with their left index finger when the central arrow (regardless of condition) faced to the left and the right tab button with their right index finger when the central arrow faced to the right. Participants were encouraged to respond as quick as possible, being accurate. A total of 120 trials were randomly presented (60 congruent and 60 incongruent trials) in each task. Each task lasted for 6 minutes approximately without breaks.

### EEG recording and analysis

EEG data were recorded at 1000 Hz using a 30-channel actiCHamp System (Brain Products GmbH, Munich, Germany) with active electrodes positioned according to the 10-20 EEG International System and referenced to the Cz electrode. The cap was adapted to individual head size, and each electrode was filled with Signa Electro-Gel (Parker Laboratories, Fairfield, NJ). Participants were instructed to avoid body movements as much as possible, and to keep their gaze on the center of the screen during the exercise. Electrode impedances were kept below 10 kΩ. EEG preprocessing was conducted using custom Matlab scripts and the EEGLAB (Delorme & Makeig, 2004) and Fieldtrip (Oostenveld et al., 2011) Matlab toolboxes. EEG data were resampled at 500 Hz, bandpass filtered offline from 1 and 40 Hz to remove signal drifts and line noise, and re-referenced to a common average reference. Horizontal electrooculograms (EOG) were recorded by bipolar external electrodes for the offline detection of ocular artifacts. The potential influence of electromyographic (EMG) activity in the EEG signal was minimized by using the available EEGLAB routines (Delorme & Makeig, 2004). Independent component analysis was used to detect and remove EEG components reflecting eye blinks (Hoffmann and Falkenstein, 2008). Abnormal spectra epochs which spectral power deviated from the mean by +/-50 dB in the 0-2 Hz frequency window (useful for catching eye movements) and by +25 or −100 dB in the 20-40 Hz frequency window (useful for detecting muscle activity) were rejected. On average, 5.1% of epochs per participant were rejected.

*Spectral power analysis*. Processed EEG data from each experimental period (Resting 1, Warm-up, Exercise, Cool Down, Flanker Task 1, Resting 2, Flanker Task 2) were subsequently segmented to 1-s epochs. The spectral decomposition of each epoch was computed using Fast Fourier Transformation (FFT) applying a symmetric Hamming window and the obtained power values were averaged across experimental periods.

*Event-Related Spectral Perturbation (ERSP) analysis.* Task-evoked spectral EEG activity was assessed by computing ERSP in epochs extending from −500 ms to 500 ms time-locked to stimulus onset for frequencies between 4 and 40 Hz. Spectral decomposition was performed using sinusoidal wavelets with 3 cycles at the lowest frequency and increasing by a factor of 0.8 with increasing frequency. Power values were normalized with respect to a ‒300 ms to 0 ms pre-stimulus baseline and transformed into the decibel scale.

## Statistical analysis

A stepwise, cluster-based, non-parametric permutation test approach (Maris & Oostenveld, 2007) was used to examine the spectral power main effect of session (moderate-to-high intensity, light intensity), separately at each period (resting 1, warm-up, exercise, cool down, task 1, resting 2, task 2) without prior assumptions on any frequency range or brain area of interest. We performed a t-test for dependent samples on all individual electrodes and frequencies pairs (30 channels × 40 frequencies), clustering samples with t-values that exceeded a threshold (p < 0.05) based on spatial and spectral adjacency. The significance of clusters was defined using 5000 permutations (see Ciria et al., 2017, for a similar approach).

For the ERSP analysis, we first tested the main effect of task condition (congruent, incongruent) separately at each flanker task (task 1, task 2) by applying the cluster-based permutation test. Subsequently, we analysed the ERSP main effect of session (moderate-to-high intensity, light intensity) using the congruency effect as dependent variable. The congruency effect was calculated through the subtraction of the two task conditions to yield the difference in ERSP activity between incongruent and congruent trials (cf. Fan et al., 2005). The ERSP main effect of session was separately calculated for each flanker task (task 1, task 2) by applying the cluster-based permutation test. Note that the EEG frequency spectrum was grouped into four frequency bands in order to reduce the possibility that the type II error rate was inflated by multiple comparisons correction: Theta (4‒8 Hz), Alpha (8‒14 Hz), lower Beta (14‒20 Hz) and upper Beta (20‒40 Hz). Additionally, the time window of interest was restricted to the first 500 ms after stimulus onset in order to avoid an overlap with behavioural responses based on average reaction time (RT).

The behavioural data analyses were performed both for RTs and accuracy (ACC) at each flanker task using statistical non-parametric permutation tests with a Monte Carlo approach (Ernst, 2004; Pesarin & Salmaso, 2010). First of all, the significant main effect of task condition (congruent, incongruent) was tested separately for each flanker tasks, with RT and ACC as dependent variables. Afterwards, we used the congruency effect (i.e. the subtraction of the two task conditions: incongruent vs congruent) as dependent variable with the within-participants factor of session (moderate-to-high intensity, light intensity) separately for each of the two flanker tasks.

## Results

### Spectral power analysis

The analysis of tonic spectral power showed a significant main effect of session for the exercise period (all *ps* < .01). Two statistically significant positive clusters (frequency-localization) were found: one global cluster (30 electrodes) in low frequencies (1-3 Hz), *p* = .009, and one centro-occipital cluster (16 electrodes) in fast frequencies (10-24 Hz), *p* = .006. The analysis revealed an overall increase in the power of frequencies during the moderate-to-high intensity exercise period in comparison to light intensity (see Fig. 2). There were no statistically significant between-session differences in any of the other periods (all cluster *ps* ≥ .1).

**Figure 2.**
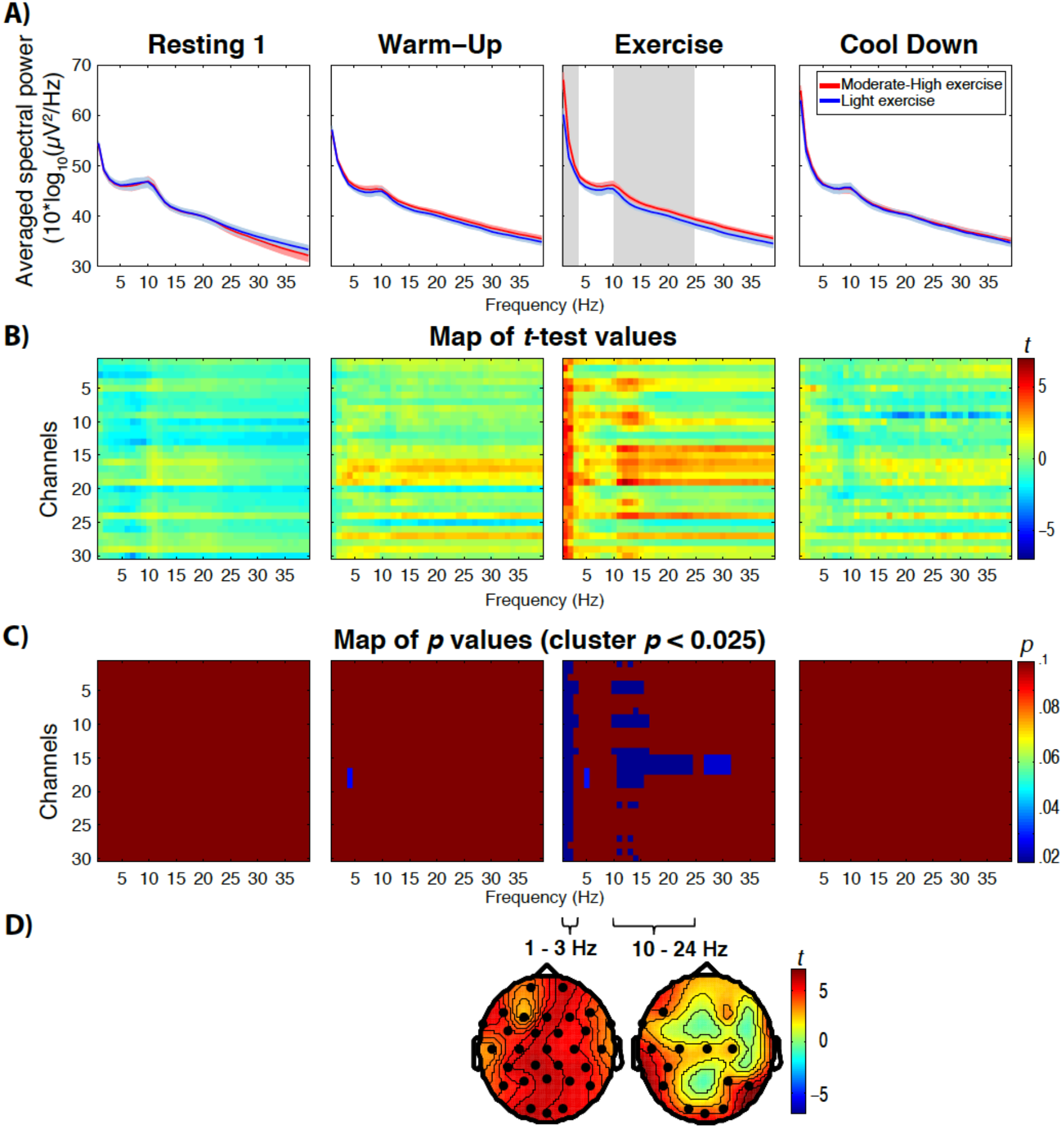
Differences in brain power spectrum as a function of exercise intensity. (A) Differences in the averaged EEG power spectrum across subjects between moderate-to-high intensity (red lines) and light intensity (blue lines) exercise at resting 1, warm-up, exercise and cool down. Red and blue shaded areas represent 95% confidence intervals. Statistically significant differences are marked by grey area. (B) Parametric paired t-test maps comparing the relative power across frequency bands (x-axes) and channels (y-axes) during moderate-to-high intensity and light intensity exercise at resting 1, warm-up, exercise and cool down (blue: decreases; red: increases). (C) Each image illustrates the statistical significance (*p* values) of the *t*-maps depicting only the significant clusters with *p* < 0.025. (D) Topographies depict t-test distribution in all electrodes, showing the spatial characteristics of the increase in power of low frequencies across the whole surface localization during moderate-to-high exercise, and the increase in high frequencies in centro-occipital areas during moderate-to-high exercise. Note that the analysis of the other periods did not yield significant between-intensity differences.

The differences of brain power spectrum as a function of exercise intensity could have been due to an increase or decrease of EEG spectral power with respect to the resting state. To address this issue, we analyzed the difference of brain spectral power during exercise with respect to the resting 1 period, separately for each exercise intensity session. The cluster-based analysis of tonic spectral power showed a significant main effect of period (Resting 1 vs. Exercise) for the moderate-to-high intensity exercise (all *ps* < .025) with two positive clusters: one global cluster (30 electrodes) in low frequencies (1-5 Hz), *p* = .01, and one tempo-occipital cluster (17 electrodes) in fast frequencies (12-39 Hz), *p* = .002. The analysis revealed an overall increase in the power of low and fast frequencies during the moderate-to-high intensity exercise period in comparison to the resting 1 period (see Fig. 3). Similarly, the analysis of the light intensity exercise showed a significant main effect of period (all *ps* < .025) with two positive clusters: one global cluster (30 electrodes) in low frequencies (1-3 Hz), *p* = .023, and one tempo-occipital cluster (11 electrodes) in fast frequencies (13-39 Hz), *p* = .017. The analysis also revealed a significant negative cluster at central locations (17 electrodes) in frequencies between 5 and 26 Hz, *p* = .010. The analysis showed an overall increase in the power of frequencies between 1 and 3 Hz, and 13 and 39 Hz during the light intensity exercise compared with the resting 1 period, parallel with a lower power between 5 and 26 Hz in central electrodes (see Fig. 3).

**Figure 3.**
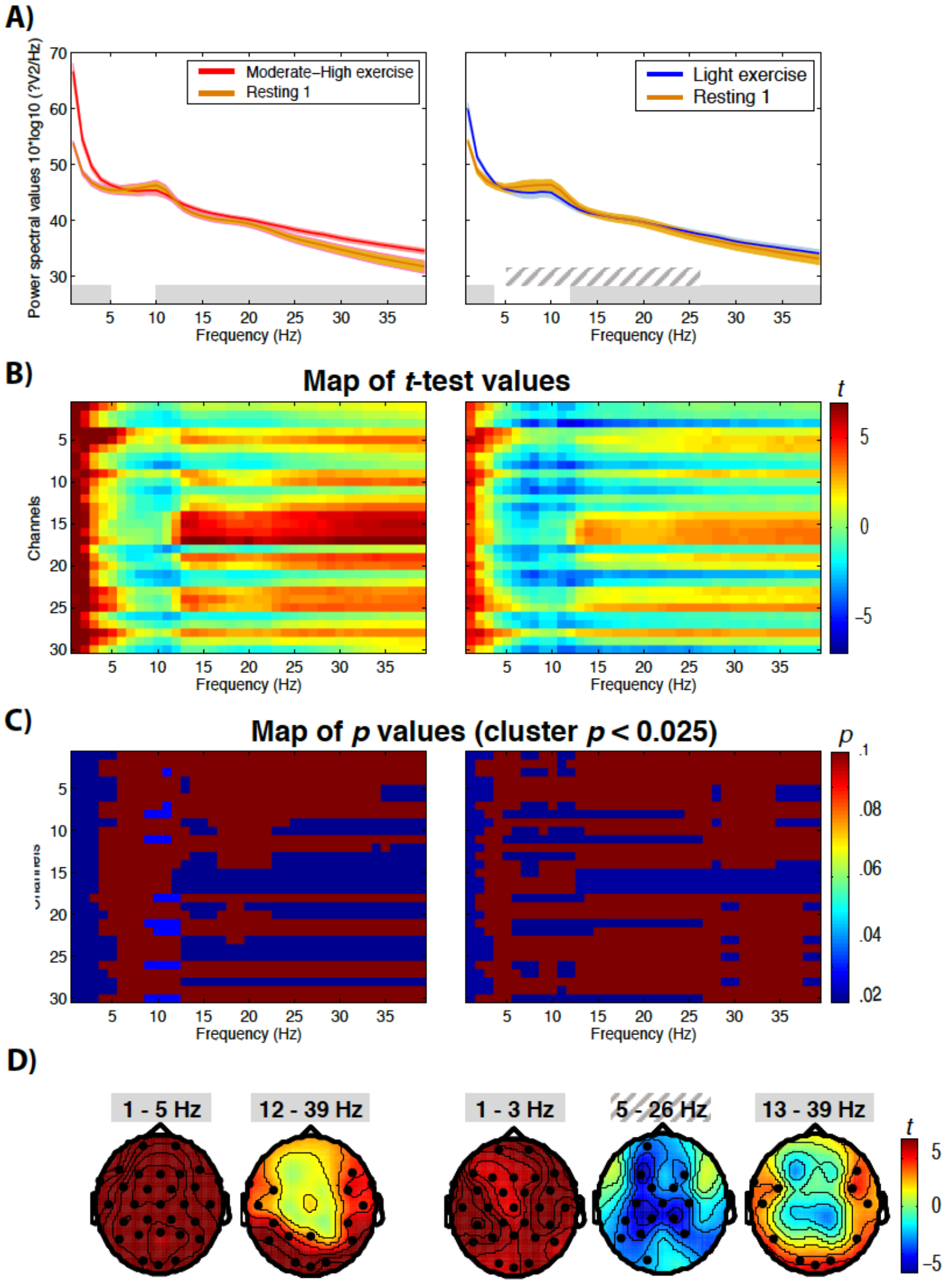
EEG spectral power as a function of exercise intensity with respect to the first resting period. (A) Left panel represent the difference in the averaged EEG power spectrum across subjects between moderate-to-high intensity (red lines) and resting 1 (yellow lines). Right panel shows the averaged EEG power spectrum difference between light intensity (blue lines) and resting 1 (yellow lines). Red, blue and yellow shaded areas represent 95% confidence intervals. Grey areas represent significant positive clusters and dashed grey area represents significant negative cluster. (B) Parametric paired t-test maps comparing the relative power across frequency bands (x-axes) and channels (y-axes) during exercise periods compared to the resting 1 (blue: decreases; red: increases). (C) Each image illustrates the statistical significance (*p* values) of the *t*-maps depicting only the significant clusters with *p* < 0.025. (D) Topographies depict t-test distribution in all electrodes, showing the spatial characteristics of the increase in power of low frequencies across the whole surface localization and the increase in high frequencies in parieto-occipital areas during both exercise intensities with respect to the resting state. The grey dashed frequency range represents the power decrease of frequencies between 5 and 26 Hz at central locations during light intensity exercise compared with resting state.

### ERSP analysis

The analysis of ERSP activity (see Fig. 4) revealed a significant main effect of task condition (incongruent vs congruent) for the flanker task 1 (all *ps* < .001). A statistically significant globally-located positive cluster (23 electrodes) in theta band between 300-500 ms after the onset of the stimuli, *p* < .001, and a significant cluster in the alpha frequency band composed by 24 electrodes between 260-500 ms, *p* < .001, were found. The analysis of task 2 showed a similar main effect of task condition (all *ps* < .001). Two positive clusters were found, one in the theta frequency band (24 electrodes) between 300-500 ms, *p* < .001, and another one in the alpha band with 21 electrodes between 270-440 ms, *p* < .001. The task condition analysis revealed a higher increase in the power of theta frequency band paralleled by a lower power suppression of alpha frequency band after the onset of incongruent trials compared to the congruent trials in both flanker tasks. The analysis of the congruency effect as a function of exercise intensity did not yield any significant difference either in the first flanker task or the second flanker task (all *ps* > .05).

**Figure 4.**
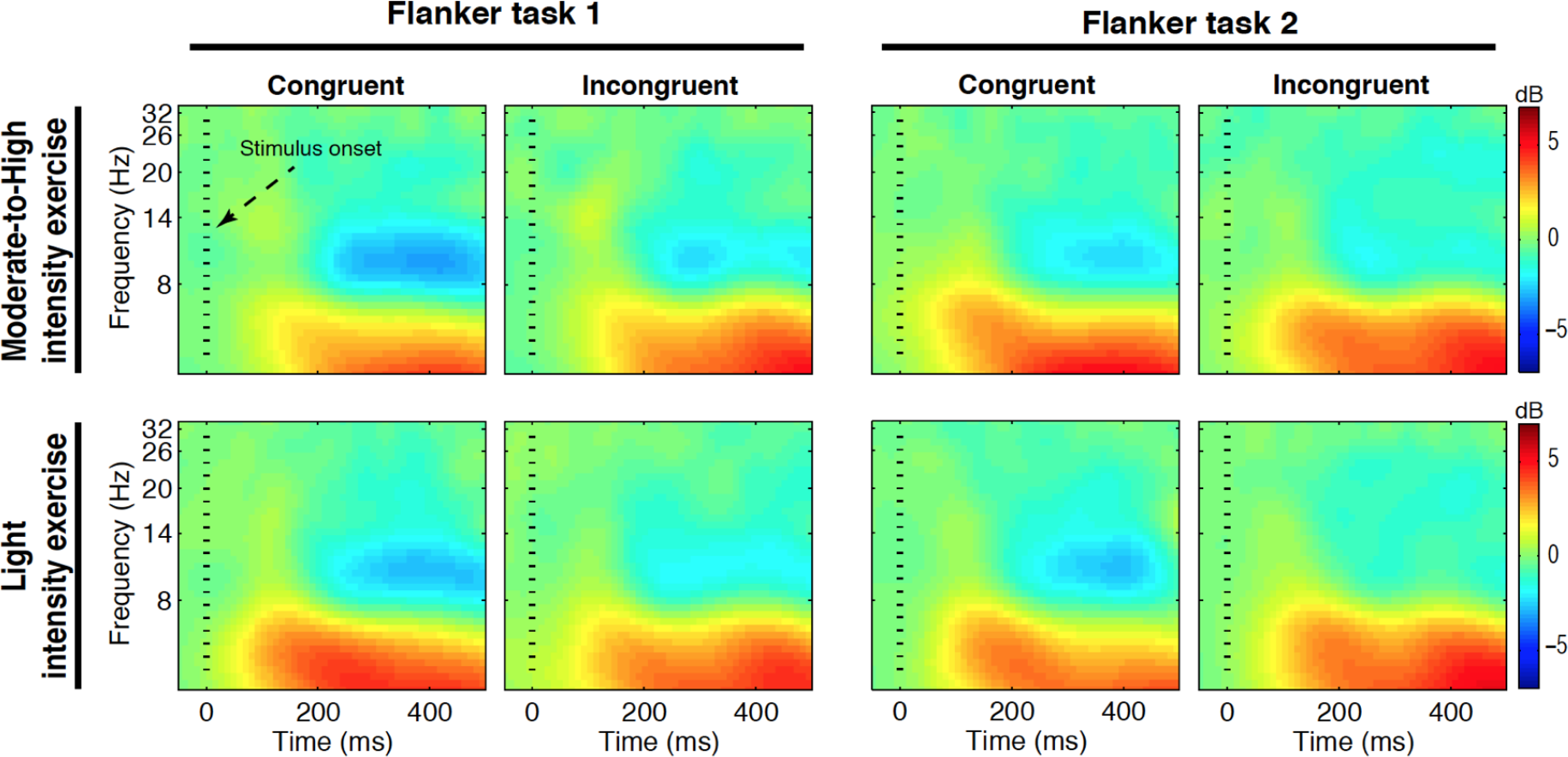
Event-related spectral perturbation of flanker tasks. Event-locked spectral power averaged at Cz channel for moderate-to-high intensity (first row) and light intensity (second row) exercise for congruent and incongruent stimuli and each task (Flanker task 1 and Flanker task 2). Each panel illustrates time-frequency power across time (x-axes) and frequency (y-axes) during moderate-intensity and light-intensity exercise (blue: decreases; red: increases).

### Behavioural Performance

Nonparametric permutation tests showed significant differences between task conditions (congruent, incongruent) for RTs and ACC in both flanker tasks (all *ps* < .001). Data revealed higher ACC and faster RTs for congruent trials with respect to the incongruent trials (see Table 1). The congruency mean RT effect was the same in the moderate-to-high intensity condition than in the light intensity condition in flanker task 1 (117 ms) and nearly the same in flanker task 2 (107 and 105 ms, for the moderate-to-high and light intensity conditions, respectively). Something similar was evident for the ACC dependent measure (see Table 1). Not surprisingly, the analysis of congruency effect for RTs and ACC with the within-participants factor of session (moderate-to-high intensity, light intensity) did not reveal statistically significant differences in any of the two flanker tasks (all *ps* > .05).

**Table 1.**
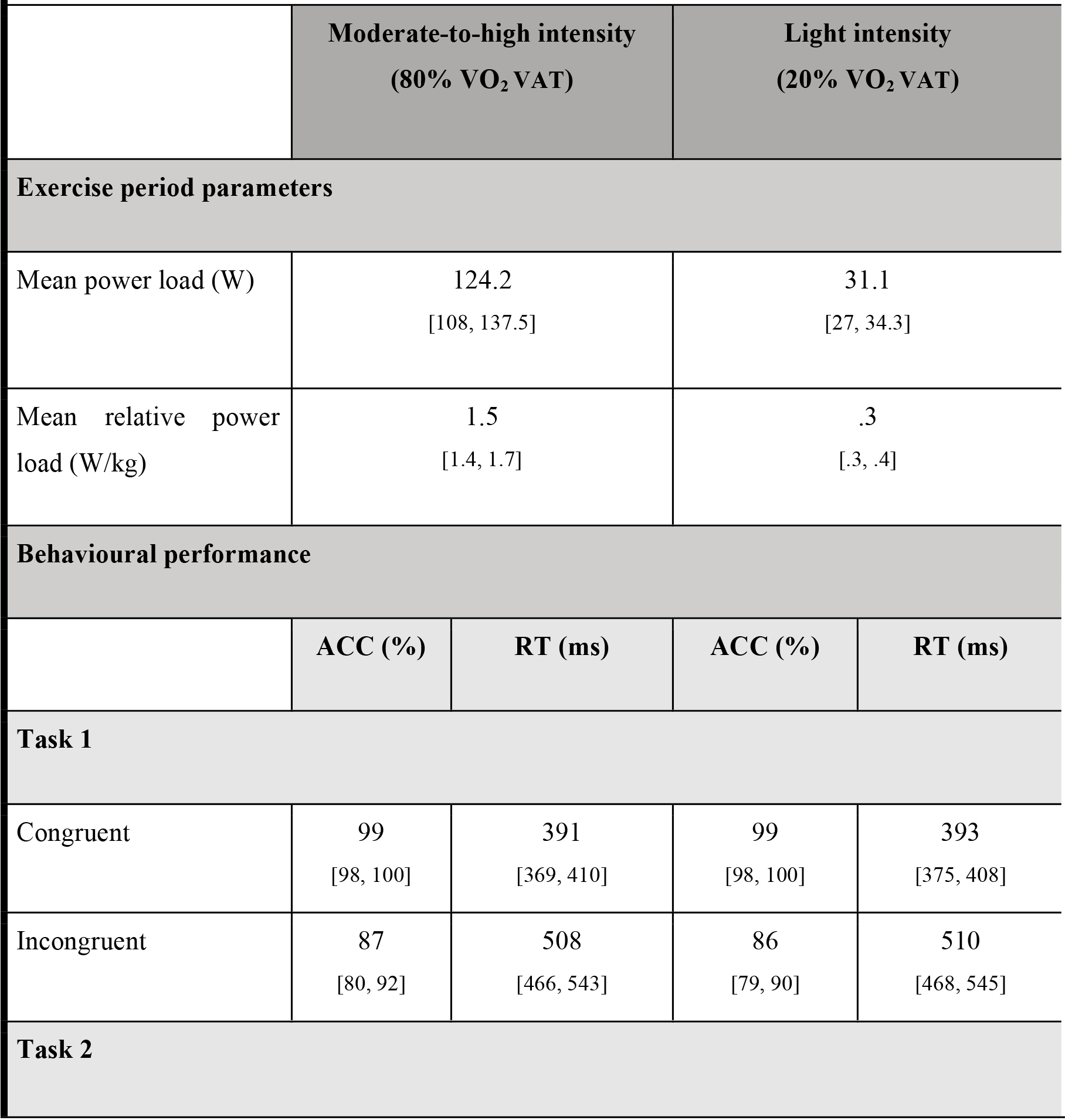
Mean and 95% confidence intervals of descriptive exercise-intensity parameters and behavioural performance for the moderate-to-high intensity and low intensity sessions.

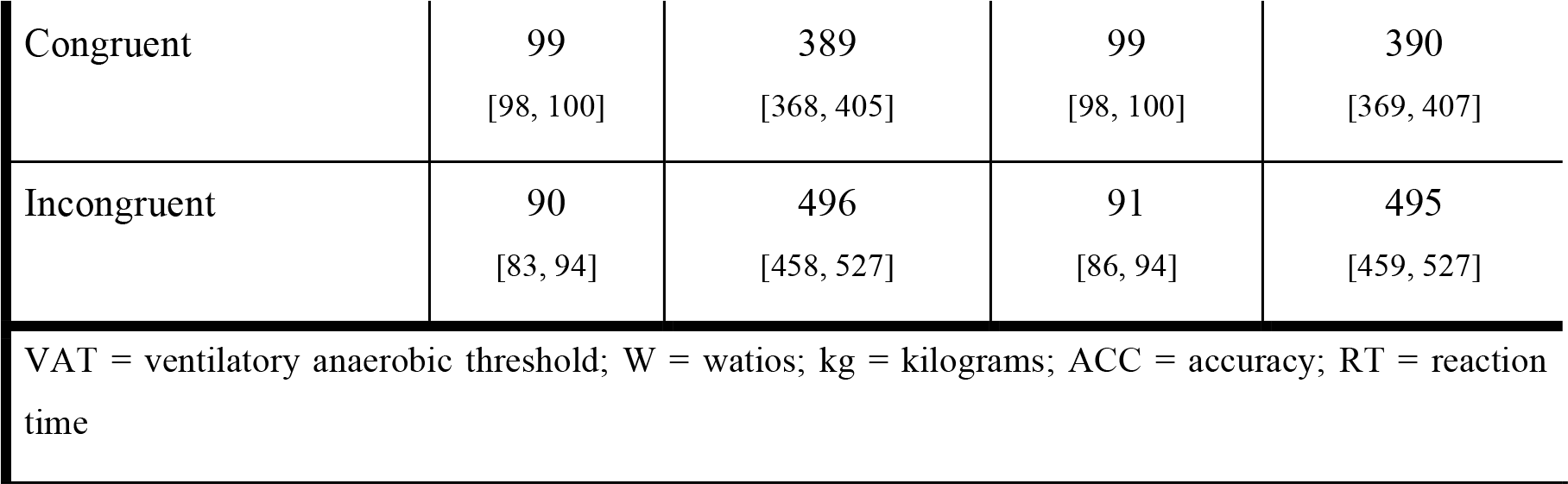

## Discussion

In the present study, we investigated the oscillatory brain activity during and following an acute bout of exercise in a group of healthy young adults as well as the impact of exercise on inhibitory control. To this end, two sessions of aerobic (cycling) exercise (i.e. moderate-to-high intensity exercise and light intensity exercise) were compared in terms of steady state EEG spectral activity. We also measured behavioural performance together with the transient (event-related) oscillatory activity during two flanker tasks (separated by a resting period) performed after the bout of acute exercise. Moderate-to-high intensity exercise, as well as light intensity exercise, induced an overall increase in the steady state oscillatory activity with respect to the resting state. This power increase was higher during the moderate-to-high intensity exercise compared to the light intensity exercise. Interestingly, the exercise-induced increase in oscillatory brain activity returned to resting levels immediately after the end of the exercise. Crucially, and in sharp contrast with previous reports (Chang et al., 2015; Verburgh et al., 2014; Chang et al., 2012; Lambourne & Tomporowski, 2010), neither the transient (event-related) oscillatory activity, nor behavioral performance during the flanker tasks following exercise showed significant between-intensity differences.

The overall power increase of the entire frequency spectrum during moderate-to-high intensity exercise with respect to light intensity is in line with previous research (Ciria et al., 2017; Crabbe & Dishman, 2004). The present results empirically confirm the absence of a selective effect of acute exercise on the alpha frequency range in anterior sites which had been taken as a potential neural mechanism underlying the beneficial effects of acute exercise on mood (Lattari et al., 2014; Boutcher, 1993; Petruzzello et al., 1991) and cognitive function (Chang et al., 2015; Dietrich, 2006). Our findings point to a generalized arousal effect of exercise that seems to influence brain oscillatory activity in several frequencies and locations. However, recent findings (Ciria et al., 2017; Ludyga, Gronwald & Hottenrott, 2016; Erickson, Hillman & Kramer, 2015) suggest that the effect of acute exercise over cognition cannot be explained as a mere overall increase of oscillatory brain activity. Instead it seems more plausible that exercise induces an efficient pattern of brain functioning, which in turn may result in improved cognitive performance.

Notably, between-intensity differences in slow frequencies were found across the entire scalp map, while differences in faster frequencies emerged from parieto-occipital locations, supporting the results reported by Ciria et al. (2017). Further, both exercise sessions elicited a global pattern of increased oscillatory brain activity with respect to the (first) resting period that was not specific to a concrete surface localization in slow frequencies, and localized in parieto-occipital electrode sites in faster frequencies. Nevertheless, resting was characterized by a similar power spectrum profile to the one elicited by moderate-to-high exercise in the range of 6 to 11 Hz, while resting EEG power was even higher than light intensity exercise EEG power between 5 and 26 Hz at central locations. It is important to note that participants were instructed to keep their eyes closed during the resting state period, which is known to drastically increase the power of alpha frequency band (Klimesch, 1999). This well demonstrated alpha peak effect could explain the absence of differences in the alpha frequency range during the moderate-to-high exercise with respect to the resting state. It could in turn also explain the lower power found during light intensity exercise in comparison to the resting state in medium frequencies.

Taken together, our findings indicate a generalized exercise-induced activation/arousal effect, similar to other physiological changes resulting from vigorous exercise, such as increases in core temperature, cortical blood flow, heart rate, or catecholamine concentration (McMorris & Hale, 2015). The direction and magnitude of these physiological changes depend on the intensity of the exercise, which has been highlighted as a key moderating variable to explain brain function and cognitive performance under physical exertion (González-Fernández et al., 2017; Chang et al., 2012; McMorris & Hale, 2012; Brisswalter, Collardeau & René, 2002). However, the absence of ERSP and behavioural differences after the end of the exercise as a function of exercise intensity do not support previous evidence pointing to a transient stimuli-sensitive modulation of specific brain rhythms associated with cognitive performance enhancement (Chang et al., 2015).

The contrast between the findings by Chang et al. (2015) and our own results could be explained by the fact that our moderate-to-high intensity exercise was compared to a (baseline) light intensity session (which we believe is the proper condition to control for the mere effect of pedaling), instead of a rest non-exercise condition (Chang et al., 2015).. The moment at which our participants completed the first flanker task (with respect to Chang et al.’s study) might be also seen as a factor contributing to our null result. However, this seems unlikely, for the inhibitory task was performed within the key time window after the end of the exercise where the largest exercise-induced benefits have been found (Chang et al., 2012; Lambourne & Tomporowski, 2010). Despite the fact that a null result (no difference in the magnitude of the congruency effect between the two effort conditions) does not imply the veracity of the alternative hypothesis, it seems to be clear that the time window where the largest cognitive benefits are expected should be revised to determine the duration of single-session exercise-induced effects on inhibitory control.

To conclude, the data we report here demonstrate an overall increase of oscillatory brain activity while exercising which seems to be unspecific of frequency range or brain location. Further, these results suggest that the heightened oscillatory power increase during exercise returns to resting levels immediately after the cessation of the exercise. Finally, the findings of the present study challenge the idea that inhibitory control benefits from a previous bout of moderate-to-high aerobic exercise.

## Acknowledgments

This work was supported by the “Ministerio de Economía y Competitividad” (grant number PSI2016-75956-P) to Daniel Sanabria, and a predoctoral grant from the Spanish Ministerio de Economía, Industria y Competitividad (BES-2014-069050) to Luis F. Ciria. The funders had no role in study design, data collection and analysis, decision to publish, or preparation of the manuscript. We thank to all the participants who took part in the experiment.

